# Interactive segmentation of membrane and membrane mimic densities in cryo-EM maps

**DOI:** 10.64898/2026.02.25.707988

**Authors:** Alok Bharadwaj, Lotte Veerbeek, Arjen J. Jakobi

## Abstract

Contextual density arising from biological membranes or membrane mimics such as detergent micelles or lipid nanodiscs, is ubiquitous in cryo-EM maps of integral and peripheral membrane proteins, or membrane-associated complexes, from single-particle analysis and subtomogram averaging. Here, we describe SURFER, a lightweight, GPU-accelerated extension for UCSF ChimeraX that enables rapid semantic differentiation of membrane or membrane mimic density from macromolecular signal. SURFER operates on unfiltered maps to automatically generate segmentation masks that can be flexibly applied to any aligned target map, including reference volumes for local refinement or post-processed maps optimised for visualisation and interpretation. By allowing contextual density to be selectively toggled or subtracted, SURFER enables direct interactive comparison between representations with and without membrane-related features to facilitate consistent analysis and reporting of membrane context in cryo-EM structures. An interactive graphical interface within ChimeraX supports convenient threshold adjustment, connectivity filtering, and immediate visual evaluation of the resulting segmentation.

## 1. Introduction

Cryogenic electron microscopy (cryo-EM) and electron tomography (cryo-ET) now routinely yield three-dimensional electrostatic potential maps for membrane proteins and membrane-associated complexes across a wide resolution range ^1,2^. Beyond the ordered macromolecular core, many cryo-EM structures of membrane proteins contain contextual density originating from sample preparation or the native environment, including detergent micelles ^3,4^, amphipols ^5^, lipid nanodiscs ^6,7^, membrane bilayers ^8^, and other partially occupied or conformationally heterogeneous components ^2^. Such contextual density is typically weakly ordered and dominated by low resolution signal. Although it can provide important information on the organisation of the membrane environment and the positioning of transmembrane regions, its incoherent signal frequently causes the density to fragment into discontinuous regions that complicate visualisation and segmentation using simple threshold-based approaches. Map post-processing has a strong influence on how such contextual density is represented. Conventional global sharpening ^9^ and many learned enhancement approaches ^10–13^ prioritise recovery of high-resolution contrast in well-ordered regions that often form the bulk of the macromolecular signal. While effective for enhancing macromolecular features, these approaches often further fragment low-resolution density or suppress it altogether ^14,15^. In contrast, map post-processing strategies that aim to preserve local signal statistics can retain contextual density at intensity levels comparable to those of the macromolecule ^16–18^. Building on these principles, LocScale-2.0 was introduced to improve the interpretability of cryo-EM maps through confidence-guided map optimisation while maintaining a more unbiased representation of both ordered and disordered components ^19^. As a result, membrane and membrane mimic density often appears more continuous and interpretable in LocScale-2.0 optimised maps at a common intensity threshold.

While retention of contextual density is advantageous for assessing membrane topology and macromolecular environment, it also introduces challenges during interactive map inspection and visual presentation. In particular, lipid or detergent density may obstruct the view on transmembrane regions or dominate visualisation at commonly used contour levels. When this density is brought onto a comparable intensity scale to the macromolecular core, interpretation and visualisation create a practical need for tools that enable semantic differentiation between contextual membrane signal and ordered macromolecular features. In addition to visual presentation, segmentation of membrane mimic density can be practically useful during 3D refinement. From an image formation perspective, a cryo-EM micrograph represents a tomographic projection of the specimen modulated by the point spread function of the electron microscope ^20^. When membrane proteins are embedded in a lipid bilayer, nanodisc or detergent micelle, these components can produce a strong contribution to the projection signal, which in some orientations may exceed that of the embedded protein itself. The strong low-resolution signal of detergent micelles or lipid nanodiscs, where detergent and lipid molecules are expected to adopt unrelated positions away from the immediate environment of the protein, can negatively impact iterative refinement procedures that assume a largely uniform signal, and can lead to overfitting. Although data-driven regularisation strategies such as non-uniform refinement ^21^ and local signal filtering ^22^ reduce sensitivity to disordered regions, selective masking and focused refinement workflows remain widely used in practice for restricting alignment and reconstruction to regions of interest or for removing non-target signal by subtraction ^23–27^.

Segmentation of cryo-EM density maps has traditionally relied on intensity-based thresholding ^28^, morphological watershed approaches ^29,30^, and region-based methods that combine connectivity with multi-scale smoothing or statistical merging ^31,32^. These approaches form the basis of widely used tools such as Segger and related workflows for partitioning macromolecular assemblies into compact subregions ^31^. More recent methods incorporate additional structural constraints, for example graph-based representations and symmetry information ^33^. While effective for separating well-defined macromolecular components, these methods are primarily designed to identify regions characterised by relatively sharp intensity gradients or compact connectivity. Lipid and detergent-associated density, by contrast, is typically smooth, weakly bounded, and sometimes spatially contiguous with the macromolecular envelope, making its segmentation sensitive to contour levels and connectivity criteria.

Similar challenges are well recognised in cryogenic electron tomography (cryo-ET) image analysis, where membranes are commonly segmented using dedicated methods developed for low signal-to-noise and diffuse boundaries, including tensor voting ^34^, deep learning-based voxel classification ^35–38^ and combinations of deep learning-based segmentation with parametric fitting and interactive curation ^39^. Such hybrid strategies are often required to accommodate the wide range of membrane geometries encountered in practice. Comparable variability is also observed in single-particle reconstructions of membrane proteins, where contextual density may arise from membrane mimics such as micelles, bicelles, nanodiscs, or planar bilayers and can differ substantially across datasets. Semantic differentiation of such density therefore benefits from combining prior information on membrane location with data-driven voxel classification. To aid in the analysis of contextual lipid and detergent densities in single-particle reconstructions and sub-tomogram averages, we have developed SURFER (Segmentation of Unstructured Regions and Filtering for Enhanced Representation), a lightweight framework for machine-learning-based segmentation of lipid and detergent densities. SURFER automatically identifies, segments, and optionally subtracts lipid and detergent densities corresponding to micelles, lipid bilayers, or lipid nanodiscs, thereby allowing users to focus alternately on the macromolecular core or the assembly in its membrane context (Figure 1). While originally designed as a companion to LocScale-2.0 map optimisation, the segmentation mask generation of SURFER operates on unfiltered maps and application of its segmentations mask is therefore compatible with any raw or post-processed map.

**Fig. 1.**
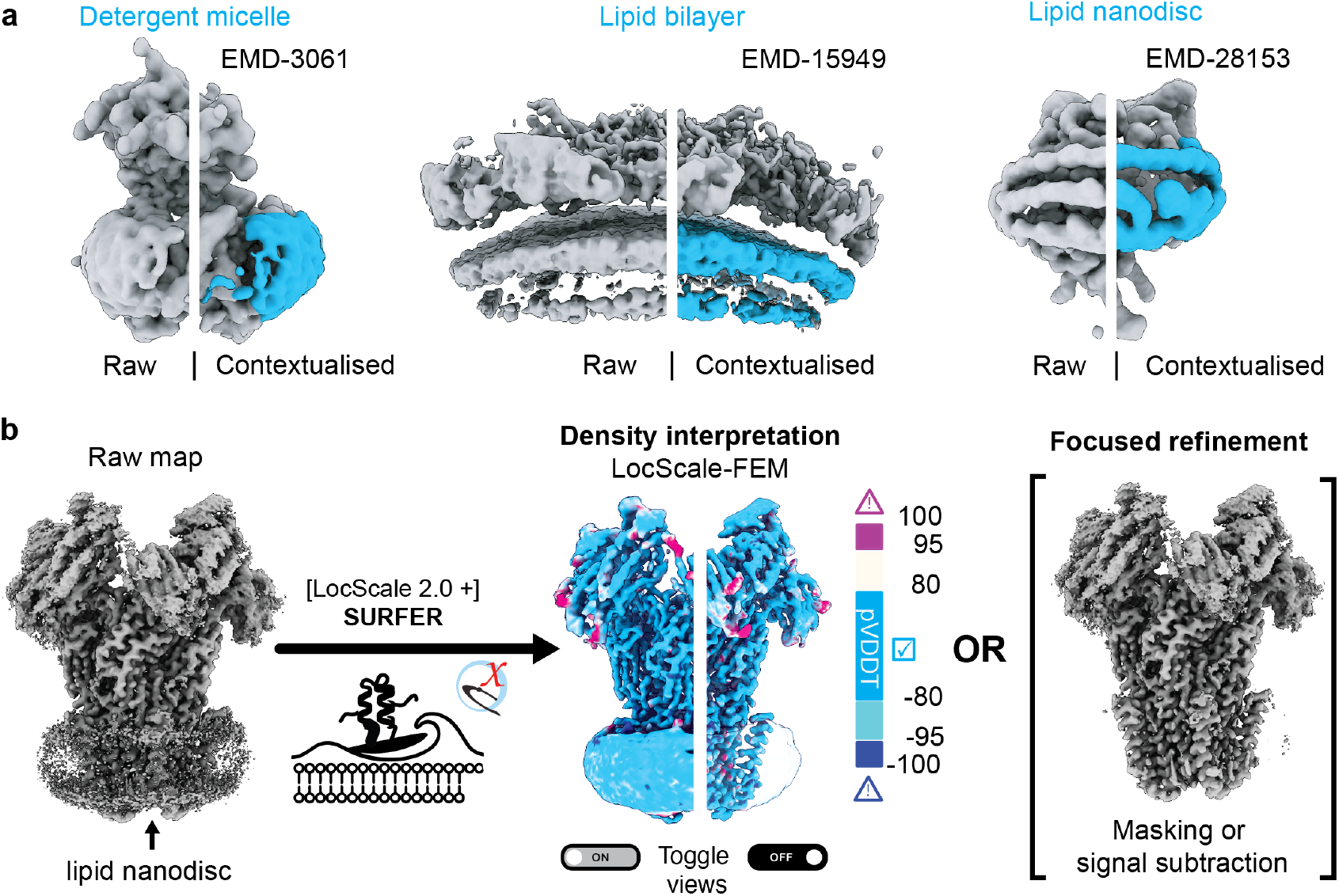
Contextualisation of lipid densities and applications of SURFER. (a) Raw and contextualised cryo-EM density maps for *γ*-secretase in amphipol A8-35 (EMDB: [EMD-3061](https://www.ebi.ac.uk/emdb/EMD-33061)), COPII coat on lipid bilayers (EMDB: EMD-15949) and the mechanosensitive channel TMEM63A in lipid nanodiscs (EMDB: EMD-28153). (b) Schematic illustration of LocScale-SURFER segmentation and substraction of lipid micelle for visualisation with LocScale 2.0-optimised maps, or for particle subtraction for 3D refinement. The example shown is the pentameric calcium-sensitive channel DeCLIC from *Desulfofustis glycolicus* reconstituted in lipid nanodiscs (EMDB: EMD-19995). The LocScale-FEM pVDDT confidence score is mapped onto the optimised map.

SURFER is implemented as a ChimeraX ^40^ bundle with both graphical user interface and command-line functionality Here, we describe the design and training of the segmentation model, evaluate its performance across a diverse set of cryo-EM density maps from EMDB-deposited membrane protein structures, and illustrate how SURFER enables contextualisation of lipid and membrane mimic densities and robust micelle subtraction in cryo-EM structures.

## 2. Results

### 2.1 Overview

Our research was motivated by a lack of accessible tools specifically optimised for segmentation of membrane or detergent micelle-related densities in three-dimensional (3D) electrostatic potential maps from cryo-EM single-particle analysis and subtomogram averaging. SURFER is a tool designed to fill this gap (Figure 1a). The method uses voxel-wise classification with a hybrid convolutional–transformer architecture to distinguish contextual density from the ordered macromolecular signal. Segmented contextual density can be then be interactively visualised or subtracted from a target map to enable direct comparison between representations that retain or suppress membrane-related features, or for the generation of masks for focused refinement (Figure 1b).

In this section, we first describe the generation of a curated training dataset and the design of the segmentation model. We then evaluate segmentation performance across a diverse set of deposited cryo-EM reconstructions and analyse the geometric variability of lipid and detergent environments captured in the training data. Finally, we illustrate practical use cases of SURFER, including interactive visualisation and subtraction of contextual density, and assess the robustness of the approach under different segmentation thresholds and filtering strategies.

### 2.2 Training dataset generation and characteristics

We assembled a labeled dataset of segmented lipid and detergent densities from curated map–model pairs of integral membrane and membrane-associated proteins derived from the Electron Microscopy Data Bank (EMDB) ^41^, restricted to entries with associated atomic models encompassing the full transmembrane region. To isolate disordered lipid and detergent density, we generated difference masks by determining the molecular envelope via statistical thresholding with false discovery rate (FDR) control ^18,19,42^ and subtracting the ordered volume occupied by the atomic coordinate model. The resulting masks contain density not explained by the atomic model, including lipid or detergent-related density together with other unmodelled structural features (Figure 2a). We then further limited this volume to membrane or membrane mimic density by predicting the planes circumscribing micelles, lipid nanodiscs or bilayer membranes around the transmembrane region using Positioning of Proteins in Membranes 3.0 (PPM 3.0), which positions membrane proteins by minimising the free energy of transfer into an anisotropic solvent model ^43,44^. While PPM 3.0 supports both planar and curved membranes, we restricted analyses here to planar geometries and reserve curved membranes for future work. Comparison of the predicted boundaries to the cryo-EM data revealed that PPM systematically underestimated lipid bilayer and micelle thickness. To correct this, we refined boundary positions by computing intensity gradients normal to the predicted planes after enhancing intensity transitions with a Sobel filter (Figure 2b). By scanning through the volume along the plane normal, the locations of maximal ascending and descending gradients can then be identified and used to define the final membrane or micelle boundaries. We applied this workflow to extract lipid and detergent densities from the averaged half maps of 254 sequence-diverse membrane protein structures deposited in the EMDB, of which 186 high-quality segmentations were retained after curation (Figure 2b, Figure S1). In most cases, these densities appear smooth because the majority of lipids or detergent molecules in membranes, micelles, or nanodiscs are mobile, resulting in averaging out of structural detail such that no individual lipid or detergent molecules can be identified. This typically means that the micelle, nanodisc or membrane components in these complexes are disordered beyond a resolution of about 10–20 Å (Figure 3a-b). The shape of these lipid or detergent structures is highly pleomorphic, depending on both the shape and symmetry of the embedded membrane protein or complex and the properties and composition of the surrounding micelle or membrane (Figure S1). We characterised this variability by measuring flatness and fragmentation levels across the micelle dataset (Figure 3c-f). The variation in flatness reflects the different physicochemical properties and molecular packing geometry of the detergent or lipid molecules in micelles, bicelles and nanodiscs. Fragmentation likely arises from low signal-to-noise ratios in the raw data. Estimating molecular boundaries when signal strength approaches noise levels is inherently challenging and can produce discontinuous, rough surfaces. While some micelles exhibit uniform fragmentation in all directions, others display anisotropy, potentially reflecting preferential protein-lipid interactions along specific orientations. Together, the flatness and fragmentation distributions shown in Figure 3c provide a compact representation of the geometric variability captured in the training dataset. For a newly analysed membrane protein structure, projecting the segmented contextual density into this feature space offers a simple way to assess whether its morphology falls within the range represented during training. Membrane mimics whose geometric descriptors lie within the main density of the training distribution are expected to be well supported by the learned model (see Section 2.3), whereas pronounced outliers may provide a practical indication that a given membrane geometry is underrepresented in the training data and could constitute a more challenging, potentially out-of-distribution case.

**Fig. 2.**
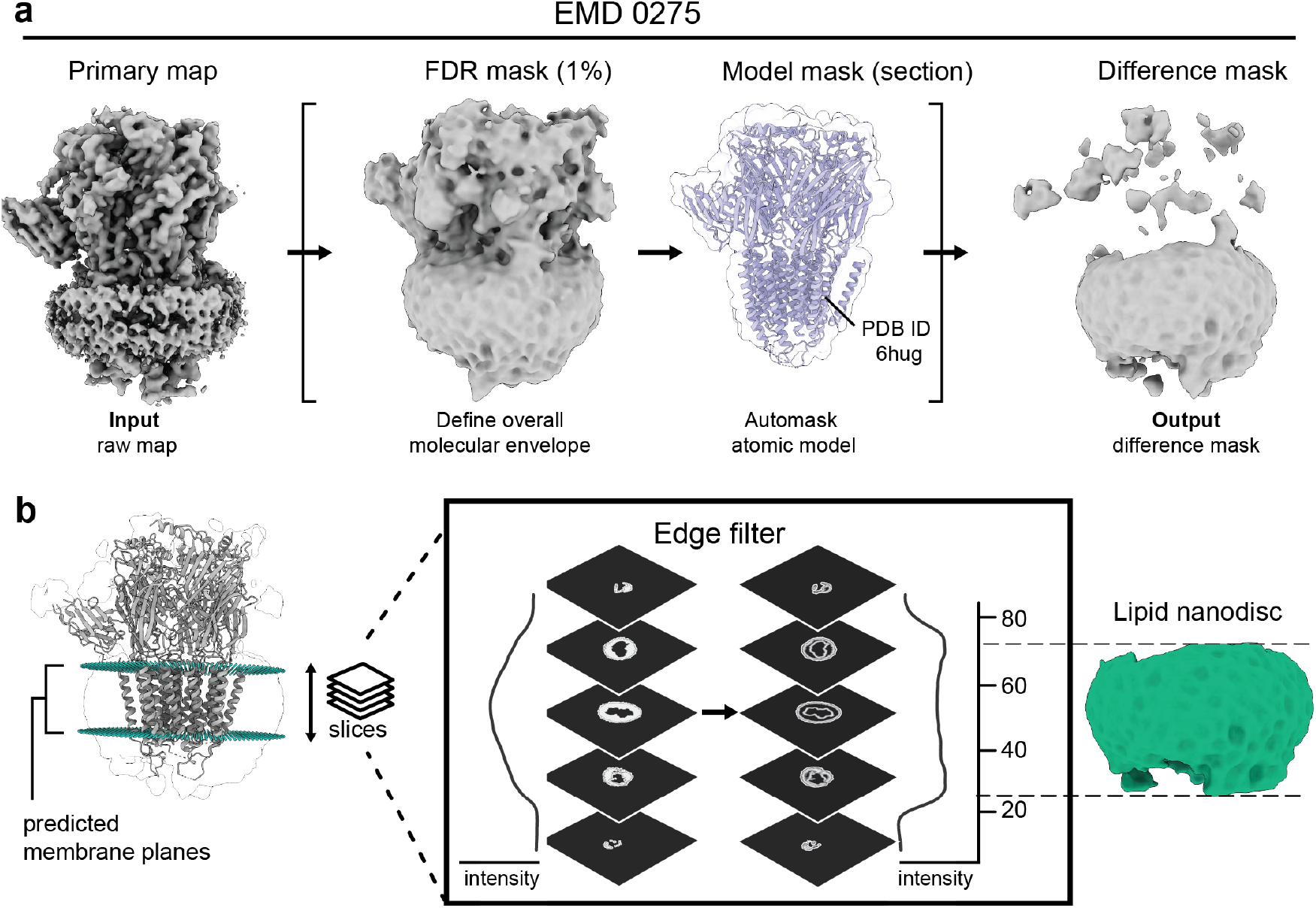
Training dataset generation for membrane/detergent segmentation. (a) Difference mask generation for isolation of lipid density regions illustrated with the α1β3γ2L GABA_A_ receptor in lipid nanodiscs (EMDB: EMD-0275). The left shows the raw average of the unfiltered half maps from which the FDR confidence mask is generated. The ordered volume is estimated from the deposited atomic model (PDB ID: 6hug) and subtracted from the FDR-enclosed volume to generate a difference mask. Note that the raw difference mask is not restricted to the lipid nanodisc but can encompass other unmodelled signal. (b) Predicted membrane planes relative to the atomic model and the raw difference mask and procedure of edge detection to refine plane positioning at the nanodisc boundary. The segmented nanodisc is shown on the far right.

**Fig. 3.**
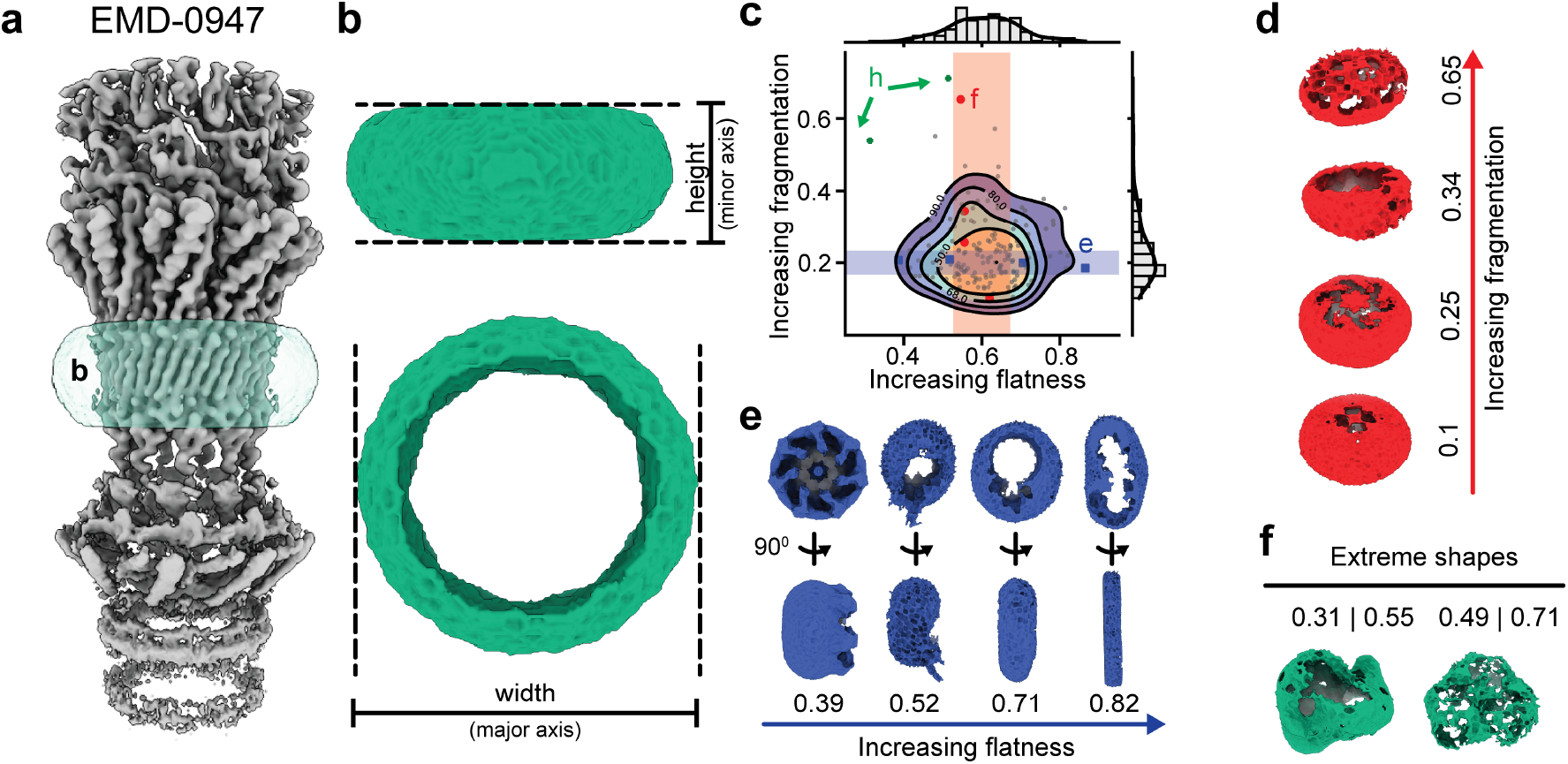
Geometric variability of membrane mimic density in the training dataset. (a-b) Geometrical properties of segmented micelle volume shown for the enterobacterial curli secretion complex CsgG-CsgF (EMDB: EMD-0947). (c) Distribution of shape features from the training dataset (N=170). Semi-transparent rectangles denote bands of largest variation along fragmentation and flatness ratios respectively. (d-e) Representative micelle densities contained within the high variability bands. (f) Out-of-distribution outliers marked by arrows in (c). Numbers indicate flatness and fragmentation ratio, respectively.

### 2.3 Segmentation model and performance

We used the annotated dataset to train a three-dimensional Swin-Conv-UNet (SCUNet) for voxel-wise segmentation of cryo-EM density maps (Figure S1). The network follows a U-Net–style encoder–decoder architecture with skip connections ^45^ and is composed of Swin-Conv blocks that combine convolutional processing with window-based vision transformer modules ^46,47^. Within each block, convolutional layers capture local density features, while transformer blocks aggregate information within fixed-size spatial windows. Although SCUNet was originally introduced for image restoration tasks ^47^, we adapt the same principle here for binary voxel-wise classification of lipid membrane or detergent micelle density versus macromolecular signal as the combination of convolutional layers and shifted window-based transformer modules enable a straighforward way for context aggregation of spatially extended density features ^48^. Details of the network architecture, training protocol, and optimisation behaviour are summarised in Table S3, Figure S2a-d and described in the Methods.

### 2.4 Threshold dependence and connectivity filtering

SURFER produces a continuous-valued voxel-wise confidence score indicating the likelihood that a given voxel belongs to membrane or membrane mimic density. To obtain a segmentation mask for visualisation or density subtraction, this output must be binarised at a chosen threshold. Because membrane mimic density typically occupies only a small fraction of the molecular volume and is dominated by weakly bounded signal, SURFER produces probabilistic outputs with rather broad confidence distributions near the class boundaries (Figure S2e). As a result, converting these probabilities into binary masks is sensitive to the chosen threshold, and the threshold that yields optimal segmentation can vary across datasets. At low segmentation thresholds, this sensitivity can lead to false-positive inclusion of low-resolution density that is not part of the membrane or membrane mimic. A good example illustrating this effect is the nanodisc-embedded transient receptor potential (TRP) ion channel NOMPC ^49^. NOMPC contains 29 ankyrin repeat (AR^1–29^) domains, of which AR^1–7^ are highly flexible and resolved only at low resolution comparable to that of the surrounding lipid nanodisc. As shown in Figure 4a and 4b, SURFER-generated segmentation masks at low thresholds include substantial off-target density corresponding to these flexible AR^1–7^ domains. Increasing the threshold progressively suppresses this off-target signal; however, rather aggressive thresholds are required before it is fully eliminated, at which point the nanodisc density itself begins to fragment and thus results in incomplete subtraction (Figure 4c). A practical way to alleviate this trade-off is to exploit the characteristic connectivity of membrane and membrane mimic density. Such density typically forms extended and spatially contiguous regions, whereas most off-target density will mainly be composed of smaller and disconnected components. SURFER therefore includes an optional connectivity filtering step in which only the largest connected component of the binarised segmentation mask is retained prior to subtraction. As illustrated in Figure 4c, including this connectivity filter can help exclude spurious off-target density while preserving the membrane mimic envelope at moderate thresholds, and thus allow clean subtraction of the lipid nanodisc.

**Fig. 4.**
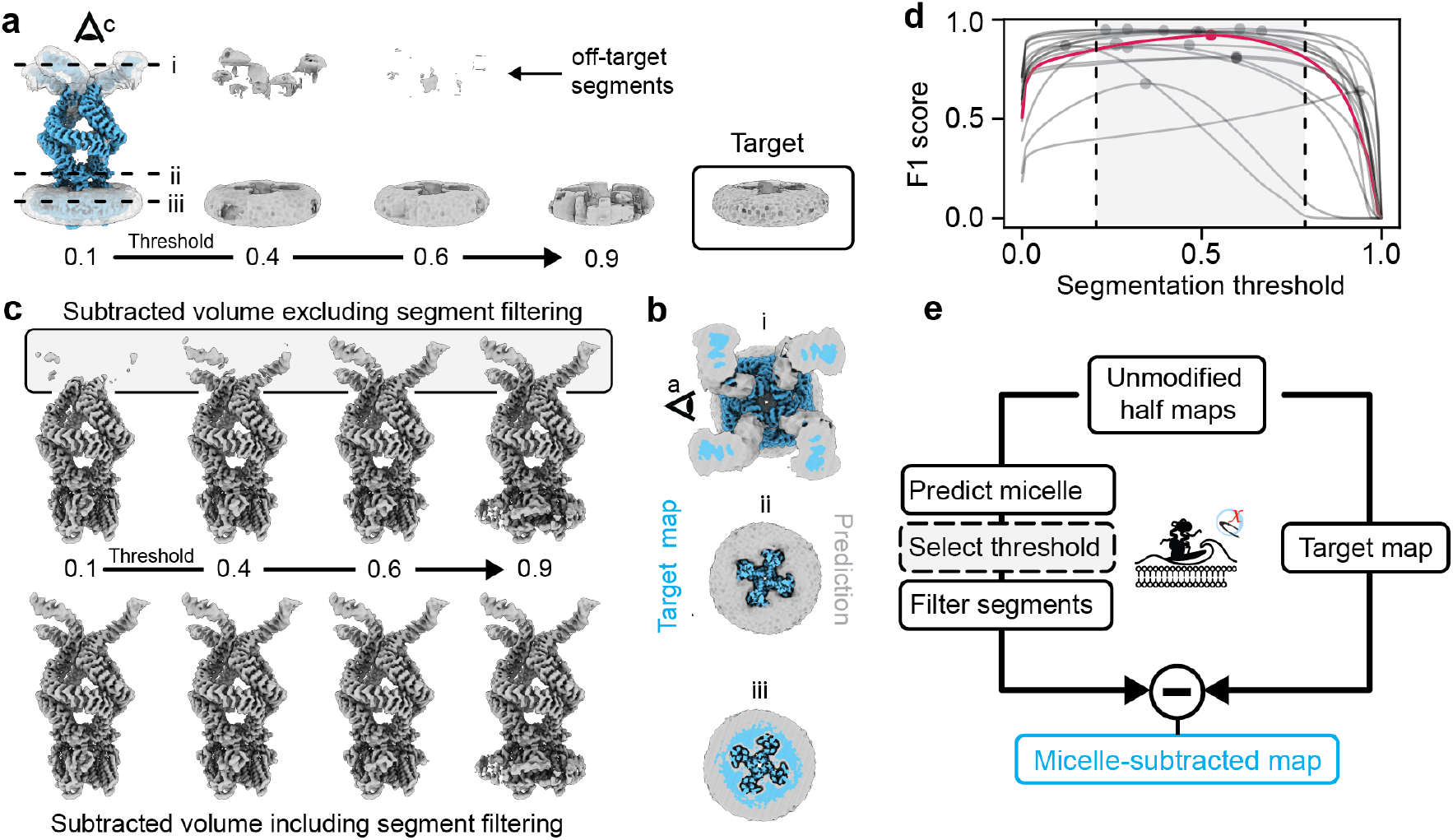
Threshold dependence and connectivity filtering. (a) Illustration of off-target segments in membrane-distal regions of nanodisc-embedded *D. melanogaster* NOMPC (EMDB: EMD-8702). Density contours for different binarisation thresholds are shown. (b) Effect of filtering based on largest connected volume on the micelle-subtracted volume. The top row shows how subtraction of the predicted segment without filtering also removes density from the off-target segments. The bottom row shows how segment filtering restricts substraction to only the micell density. (c) 2D section views at the three section planes indicated in (a) shows the overlap of the unfiltered micelle prediction with the target map. (d) F1 score as a function of binarisation threshold evaluated on the held-out test dataset. Segmentation performance peaks at an intermediate threshold, reflecting the trade-off between recall at low thresholds and precision at high thresholds. The position and sharpness of this optimum vary across reconstructions. The curve for EMD-8702 is shown in red. (e) Schematic illustration of the interactive thresholding workflow implemented in SURFER, highlighting how users can explore the effect of threshold choice on segmentation quality and apply connectivity filtering prior to density subtraction.

To more generally examine the effect of binarisation threshold on segmentation quality, we evaluated voxel-wise segmentation accuracy as a function of threshold using the F1 score, computed on the held-out test dataset for which 16 curated reference segmentations were available. The F1 score, defined as the harmonic mean of precision and recall, provides a measure of segmentation performance that is particularly informative in the presence of strong class imbalance as is the case here (Figure S2d). As shown in Figure 4d, the F1 score exhibits a pronounced dependence on the chosen segmentation threshold. For any given reconstruction, there exists a threshold at which the best balance is achieved between recovering the membrane mimic density and avoiding false-positive inclusion of unrelated low-resolution signal. At low thresholds, recall is high but precision is reduced due to inclusion of off-target low-resolution signal, leading to moderate overall performance. Increasing the threshold suppresses these false positives and improves precision, resulting in a peak F1 score at an intermediate threshold. At still higher thresholds, recall decreases as the predicted membrane mimic density becomes fragmented or incomplete, again reducing the F1 score. The location and shape of this optimum varies between reconstructions but for most reconstructions F1 scores are relatively flat across a range of segmentation thresholds centred on 0.5. This behaviour highlights two points: First, even though a generic 0.5 threshold appears well-suited as a reasonable best guess, segmentation quality can probably not be reliably optimised using a single global threshold across datasets. Second, the threshold dependence observed in the quantitative evaluation mirrors the practical challenges encountered during visual inspection, where users may have to balance removal of smooth contextual density against preservation of weak but structurally relevant features. These observations motivate the interactive threshold selection and connectivity-based filtering implemented in SURFER, which allow users to adapt the segmentation to the specific characteristics of each reconstruction rather than relying on a fixed global threshold (Figure 4e & S2f).

### 2.5 Interactive segmentation and context exploration

SURFER is deployed as an interactive plugin within the UCSF ChimeraX environment (Figure 5a). ChimeraX is increasingly used for atomic coordinate and density map display and manipulation, making it a natural platform for a user-friendly contextual segmentation workflow. The plugin provides a ChimeraX interface to the SURFER tool, enabling users to perform contextual segmentation directly during map inspection and analysis (Figure 5b). However, all SURFER functions can also be programmatically accessed via the command line if desired. The workflow comprises an automated segmentation step followed by interactive threshold refinement and application. The SURFER tools allows users to specify a pair of unfiltered, independent half maps as input. From these, SURFER computes a FDR-controlled confidence mask to define the molecular boundary and subsequently predicts detergent- or membrane-associated density within this region. The predicted contextual density can be interactively thresholded to generate a binary segmentation mask, which can then be applied to any target map aligned with the input to selectively remove micelle or membrane signal (Figure 5b). Target maps may include the unfiltered input map itself, for example, to generate masks for local 3D refinement, or post-processed maps intended for display and figure generation, such as contextualised maps produced by LocScale-2.0 FEM map optimisation.

**Fig. 5.**
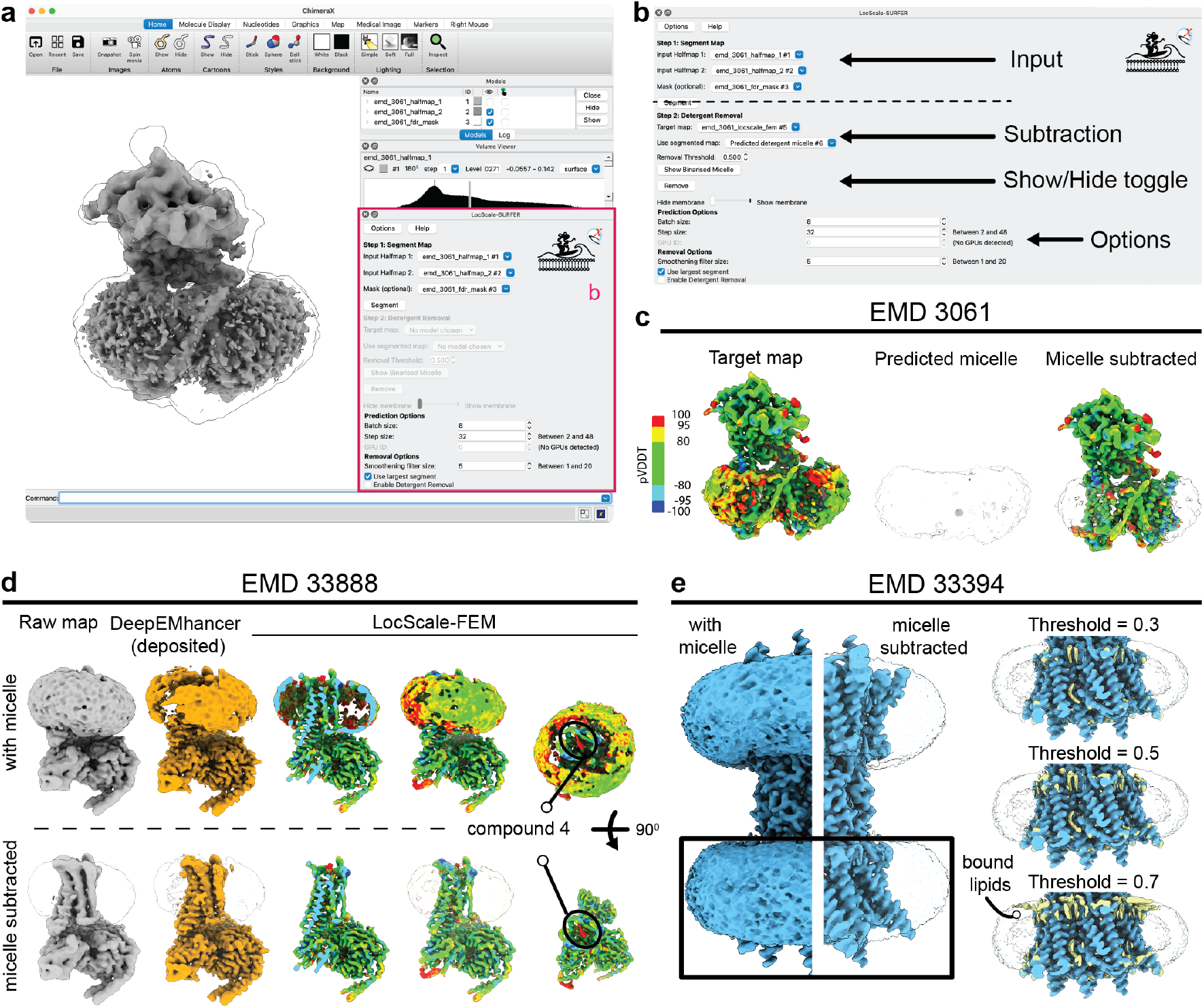
Interactive segmentation and contextual density control with SURFER in UCSF ChimeraX. (a) ChimeraX interface illustrating SURFER applied to the *γ*-secretase complex (EMDB: EMD-3061). The averaged unfiltered map is shown in grey, with the outline of the FDR-based confidence mask superposed. (b) Close-up of the SURFER tool interface. SURFER accepts a pair of unfiltered half maps as input and optionally a molecular boundary mask. Predicted micelle density can be visualised across a range of binarisation thresholds. Once a threshold is selected, the segmented micelle can be subtracted from any target map aligned to the raw input. In this example, a LocScale-FEM optimised map19 is used as the target. (c) LocScale-FEM target map, SURFER-predicted micelle density, and the corresponding micelle-subtracted map for EMD-3061. (d) Application of SURFER to the relaxin family peptide receptor 4 (EMDB: EMD-33888). Shown are the raw map, the deposited DeepEMhancer-optimised map, and the LocScale-FEM map before (top) and after (bottom) micelle subtraction. Density corresponding to the bound RXFP4 ligand is preserved int he subtracted map. (e) Application of SURFER to the connexin43 (Cx43/GJA1) gap junction intercellular channel (EMDB: EMD-33394). A LocScale-2.0 optimised target map is shown before and after micelle subtraction. Density associated with tightly bound lipids is retained in the micelle-subtracted map, although its retention depends on the chosen binarization threshold.

The ChimeraX interface allows users to toggle between representations with and without contextual density to facilitate direct visual comparison. Depending on map size, the procedure typically completes within seconds for smaller reconstructions and within approximately 10 minutes for the largest maps tested (Table 1).

**Table 1.** SURFER runtime for selected EMDB entries.

| EMDB | Box size (voxels) | Pixel size (Å) | Box size (Å) | Volume ($\times 10^6$ Å<sup>3</sup>) | Time |
| --- | --- | --- | --- | --- | --- |
| 30017 | 240 | 1.03 | 247.2 | 15.1 | 36–38 s |
| 3061 | 180 | 1.40 | 252.0 | 16.0 | 30–40 s |
| 8702 | 400 | 1.20 | 480.0 | 110.6 | 3–4 min |
| 13234† | 336 | 1.70 | 571.2 | 186.3 | 4–5 min |
| 19999† | 460 | 1.63 | 749.8 | 421.5 | 10–12 min |

† Map does not contain micelle density; used to quantify runtime only.

Figure 5c illustrates this workflow for the human γ-secretase complex (EMDB: EMD-3061) ^26^. In this example, the target map is a LocScale-2.0 feature-enhanced map (LocScale-FEM), in which ordered macromolecular regions and more poorly resolved contextual density including the detergent micelle are represented on a comparable intensity scale. Application of SURFER segmentation to the LocScale-FEM map enables selective removal of the surrounding micelle while preserving the protein-associated transmembrane features that benefit from LocScale-2.0 optimisation such as the TMD2 (Figure 5c). We also tested the ability of the segmentation workflow in distinguishing smooth contextual density arising from mobile detergent molecules or lipids from density corresponding to ordered lipids or small-molecule ligands at the protein–micelle interface. Figure 5d shows the result for the relaxin family peptide receptor 4 (RXFP4; EMDB: EMD-33888), which contains a weakly bound amidrazone scaffold compound located at the transmembrane region ^19,50^. We performed micelle subtraction on the raw reconstruction, the author-deposited DeepEMhancer-optimised map, and a LocScale-FEM map of RXFP4. Comparison of the maps before and after micelle subtraction shows that, in this case, SURFER effectively removes the surrounding micelle while preserving the ligand density at the automatically determined segmentation threshold. There are, however, cases in which the segmentation mask at the default threshold may encroach upon less well-ordered density in transmembrane regions. Figure 5e illustrates application of SURFER to the hexameric connexin43 (Cx43/GJA1) gap junction intercellular channel (EMDB: EMD-33394) in lipid nanodiscs, again using a LocScale-2.0–optimised target map ^19,51^. The Cx43/GJA1 structure of EMD-33394 (PDB: 7xqf) contains 24 modelled copies of the cholesterol ester cholesteryl hemisuccinate (CHS) and 12 copies of the phospholipid 1-palmitoyl-2-oleoyl-phosphatidylethanolamine (POPE) tightly associated with the transmembrane domains of both hemichannels. When binarising the segmentation mask at the default threshold of 0.5 or lower, in this case, the retention of ordered lipid features depends on the chosen binarisation threshold for the segmentation mask, reflecting their partial spatial overlap with the segmented contextual density. It thus appears that interactive threshold adjustment can be necessary to balance suppresion of smooth contextual density and retention of structured, protein-associated lipid density.

### 2.6 Generalisation across membrane mimics

To assess the ability of SURFER to segment contextual density across a broad range of membrane environments, we investigated a set of 20 cryo-EM structures of the bacterial ATP-binding cassette transporter MsbA determined in 12 distinct membrane mimetic systems (EMDB: EMD-50774–EMD-50794). This dataset provides a controlled benchmark in which in which the macromolecular core adopts a limited set of conformations, while the surrounding contextual density varies substantially in geometry, intensity and other characteristics. The series includes classical detergents (DDM, LMNG, GDN, Triton, UDM), nanodiscs assembled with different membrane scaffold proteins (MSP) variants, peptidiscs, and amphipols ^4^. These membrane mimics produce contextual density ranging from compact, approximately spherical micelles to flattened nanodiscs and irregular peptidisc belts (Figure 6a). We applied SURFER segmentation to all 20 reconstructions using the default binarisation threshold and subtracted the predicted lipid or detergent density from each map (Figure 6a). Across the set of 20 maps, SURFER consistently identified the dominant detergent or lipid envelope surrounding MsbA despite pronounced differences in micelle shape and thickness. Differences in segmentation quality appear primarily associated with variations in signal continuity and resolution of the membrane mimic density rather than systematic misclassification. Subtraction of the predicted contextual density preserved the structured transmembrane helices and cytosolic domains of MsbA across the series. In several nanodisc and amphipol reconstructions, density corresponding to membrane scaffold proteins or polymer belts remained partially visible (arrows in Figure 6a). These results indicate that SURFER generalises across a broad range of experimentally distinct membrane mimetic systems without mimic-specific parameter adjustment and captures shared geometric and intensity features of contextual lipid and detergent density.

**Fig. 6.**
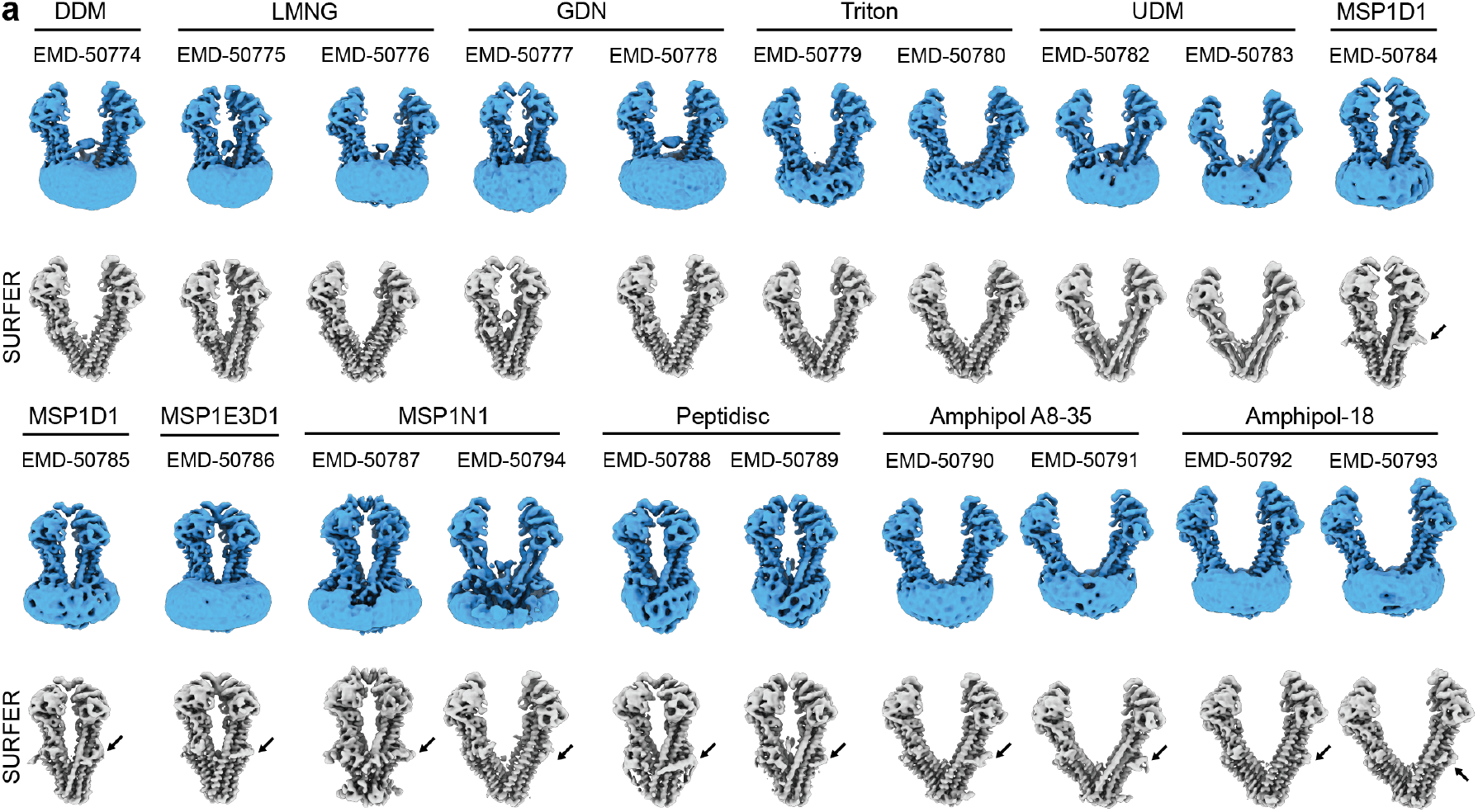
Segmentation and subtraction of diverse membrane mimics for MsbA. (a) LocScale-FEM maps of MsbA solved in 12 different membrane mimics (EMDB: EMD-50774 through EMD-50794) before (blue) and after (grey) segmentation and micelle subtraction with SURFER. Arrows indicate retained density of membrane scaffold proteins, petidiscs and amphipols.

## Discussion and conclusion

Automated segmentation of contextual membrane and membrane mimic density in cryo-EM maps poses challenges distinc from from those encountered when segmenting density of ordered macromolecular components.

Primarily inspired by LocScale-2.0^19^, which optimises map representations to emphasise contextual density, SURFER was built to provide automated segmentation of membrane-related signal together with interactive control over its visualisation. While not explicitly tested for this purpose, the procedure may in principle facilitate mask generation for membrane subtraction or focused refinement. This could be advantageous for small integral membrane proteins, where dominant low-resolution micelle or nanodisc signal may influence a lignment. However, caution is warranted as subtracting signal from particle images or reconstructions risks removal of structurally relevant signal. While the observed robustness of SURFER segmentation is encouraging in this respect, systematic evaluation of its suitability for subtraction-based and focused refinement strategies remains the subject of future work.

The results presented here show that SURFER can reliably identify membrane mimetic density across a wide range of membrane protein structures and membrane mimic geometries. In many cases, default segmentation parameters appear sufficient t o a ccurately separate and subtract contextual density while preserving ordered transmembrane regions. In more challenging situations, interactive adjustment of the binarisation threshold may be required to balance removal of smooth contextual signal against retention of structured features. The segmentation and subtraction procedure typically completes within a few minutes on a conventional laptop, making it well suited for interactive use.

The integration of SURFER into the UCSF ChimeraX ^40^ environment allows users to directly assess the plausibility of the segmentation, compare maps with and without contextual density, and generate masks for downstream applications without leaving the visualisation environment. SURFER appears to generalise reasonably well across different membrane mimics but is less accurate for membrane geometries that are underrepresented in the training data, such as strongly curved lipid bilayers. In such cases, segmentation quality is reduced and greater user intervention may be required. Extending the training dataset to include a broader range of membrane geometries is therefore a natural direction for future work.

Overall, SURFER provides a straightforward and accessible approach for contextual density segmentation in cryo-EM maps. By enabling selective suppression or visualisation of membrane and detergent density during interactive analysis, it complements existing map processing and visualisation tools and facilitates more objective interpretation of membrane protein structures.

## ACKNOWLEDGEMENTS

We thank Tom Goddard for useful suggestions concerning UCSF ChimeraX integration, as well as Arne Möller, Kilian Schnelle and the participants of the CCP-EM Icknield workshops for testing early versions of the method. This work was supported by the European Research Council (ERC) under the European Union’s Horizon 2020 research and innovation programme (ERC-StG-852880 to AJ) and CCP-EM. We acknowledge the use of computational resources of the DelftBlue supercomputer provided by the Delft High Performance Computing Centre.

## DATA AND CODE AVAILABILITY

SURFER is open-source software released under a BSD license and is available at https://github.io/cryotud/locscale-surfer, where detailed documentation is also provided. SURFER is also available directly from the ChimeraX toolshed at https://cxtoolshed.rbvi.ucsf.edu. The SURFER segmentation model (v0.1) can be accessed via Zenodo (DOI: https://zenodo.org/records/15488062). Scripts used for analysis and for the generation of figures shown in this manuscript can be found at https://github.com/cryotud/publications.

## Methods

### Software environment

Model training and inference were implemented in PyTorch (v2.9) with CUDA 12.6. Data processing and analysis were performed in Python (v3.11). Cryo-EM map and atomic model operations were carried out using EMmer ^18^, a Python library for cryo-EM map and model processing built on mrcfile^52^ and gemmi ^53^.

### Training dataset generation

#### Selection of map–model pairs

Training data were derived from cryo-EM reconstructions containing detergent micelles or membrane-associated density. Half-map pairs for 500 EMDB entries were initially collected. From these, 254 reconstructions containing visible lipid membranes, nanodiscs or detergent belts were identified through automated filtering and manual inspection. From this initial set, only entries with corresponding atomic models deposited in the Protein Data Bank (PDB) ^54^ and annotated membrane orientations in the OPM database ^55^ were retained. After target-mask generation and manual quality control (see below), 186 map–model pairs were retained. Of these, 144 were used for training and 26 for validation (85:15 split), while 16 were reserved as an independent test set. The full curated set is listed in Table S1 and S2.

#### Molecular envelope and difference mask generation

For each reconstruction in the curated training and validation set, a mask defining the molecular envelope was generated using false discovery rate (FDR)–controlled thresholding ^19,42^. Unfiltered half maps were averaged prior to FDR estimation. Noise statistics were estimated from cubic regions extracted from the edges of the reconstruction volume. The noise box size was set to the larger of 20^3^ voxels or 10% of the box dimension of the reconstructed volume. To obtain smooth and conservative molecular boundaries, maps were low-pass filtered to 5 Å before FDR thresholding. All masks were binarised at an FDR of 1% and used consistently throughout training and validation. Density contained in the FDR mask but not explained by the atomic model was identified by subtracting a model-derived mask from the FDR mask. Model masks were generated from simulated model maps using gemmi ^53^ by including all voxels within 3.0 Å of any atom in the deposited structure. The resulting difference mask frequently contained thin shell artefacts at protein boundaries due to minor discrepancies between model and envelope estimation. These artefacts were removed by uniform filtering followed by re-binarisation.

#### Prediction and refinement of membrane boundaries

Approximate membrane orientations were obtained from OPM ^55^. Because OPM coordinates are not aligned to deposited PDB models, all OPM structures were rigid-body aligned to their corresponding PDB entries using `gemmi`. Membranes were assumed to be planar for training data generation; curved membranes were excluded. Each membrane was represented by two parallel planes of the form

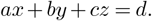

Initial plane parameters were estimated from three non-collinear points sampled from the predicted membrane surface. To refine the plane positions, edges in the difference mask were enhanced using a Sobel filter. Slices were extracted along the direction normal to the membrane planes, and the number of foreground voxels per slice was computed to generate an intensity profile. Peaks in the gradient of this profile were used to determine the refined membrane boundaries. The difference mask was subsequently cropped between the two refined planes.

### Geometric characterisation of contextual density

To quantify geometric variability across detergent and membrane-associated densities, two complementary descriptors were computed from binarised contextual volumes: a flatness ratio and a fragmentation ratio. Volumes were binarised at a threshold of 0.5 prior to feature extraction.

#### Flatness ratio

The flatness ratio provides a measure of the aspect ratio of the micelle or membrane envelope. A best-fitting ellipsoid was computed for each binarised volume using `pyradiomics` ^56^. The height of the micelle was defined as the minor principal axis of the ellipsoid and the width as the major principal axis. The flatness ratio was defined as

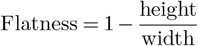

Values approaching 1.0 indicate highly flattened, bilayer-like geometries, whereas lower values correspond to more isotropic or weakly anisotropic detergent belts.

#### Fragmentation ratio

Spatial continuity of the segmented density was quantified using a surface-area-to-volume metric that describes the degree of fragmentation. Binarised volumes were converted to triangular meshes using the marching cubes algorithm ^57^. The surface area was computed as the sum of triangle areas, and the enclosed volume was obtained by tetrahedral decomposition.

The fragmentation ratio was defined as

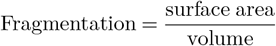

Because area and volume are computed in physical units using the map pixel size, this ratio has units of Å^-1^. Higher values indicate increased roughness or fragmentation.

#### Connectivity filtering

A binary segmentation sometimes contain multiple mutually disconnected regions, including unrelated low-resolution density. Connected-component analysis was done using label function implemented in `scikit-image` and only the largest connected component was retained. The filtered volumes were manually inspected for dataset curation. Entries for which micelle density could not be reliably isolated were excluded. This curation step yielded the final dataset used for training, validation, and testing.

### Segmentation network

#### Network architecture

SURFER employs a 3D Swin-Conv U-Net (SCUNet) ^47^. The encoder consists of three locks, each composed of a Swin-Conv module followed by strided convolution for downsampling. Within each Swin-Conv module, window-based transformer layers operate in parallel with convolutional layers and their outputs are concatenated prior to resolution reduction. A single Swin-Conv module forms the bottleneck. The decoder mirrors the encoder using transposed convolutions for up-sampling and skip connections to corresponding encoder features. Architectural details are summarised in Table S3 and Figure S2a.

#### Pre-processing and augmentation

All input maps were resampled to 1 Å/voxel and intensities were standardised to zero mean and a standard deviation of 0.1. Data augmentation was applied prior to cube extraction and included random rotations about Cartesian axes, translations up to ±10 voxels, B-factor modulation (random shifts between 0 and 400 Å^2^; applied twice per map), and Gaussian blurring corresponding to effective resolutions between 5 and 20 Å.

#### Subvolume sampling

Training samples consisted of cubic subvolumes of size 48 Å. Cubes were extracted on a regular Cartesian grid with 26 Å spacing. Grid positions were classified as signal or noise using the resampled FDR mask. All signal-containing cubes were retained. Additional noise cubes were randomly sampled to achieve a signal-to-noise cube ratio of 4:1.

#### Training parameters

The network was trained using cube-wise binary cross-entropy (BCE) loss with L1 regularisation and the Adam optimiser with default momentum parameters (β = [0.9, 0.999]). Training was performed over eight epochs. The optimal weights for the L1 regularizer, learning rate, and batch size were determined empirically using the Optuna optimization framework ^58^. The optimal values were found to be as follows: L1 weight decay of 0.0025, a learning rate of 5*×*10^−4^, and a batch size of 64.

### ChimeraX integration

SURFER is distributed as a ChimeraX bundle and can be installed via the [ChimeraX Toolshed](https://cxtoolshed.rbvi.ucsf.edu/apps/chimeraxlocscalesurfer) or from the [source repository](https://github.com/cryoTUD/locscale-surfer). In ChimeraX, the tool is accessible under *Tools → Volume Data*. The bundle includes all required metadata and trained model weights within the source distribution. The workflow comprises two stages. First, unfiltered half maps are supplied for segmentation. If preferred, a user-defined mask can also be provided. Second, the predicted segmentation is applied to any aligned target map. Users select a binarisation threshold to define the final segmentation mask, which may be optionally smoothed using a uniform filter with a default width of 5 voxels. Contextual density can be interactively toggled to compare representations with and without membrane signal.

## Supplementary material

**Table S1.**
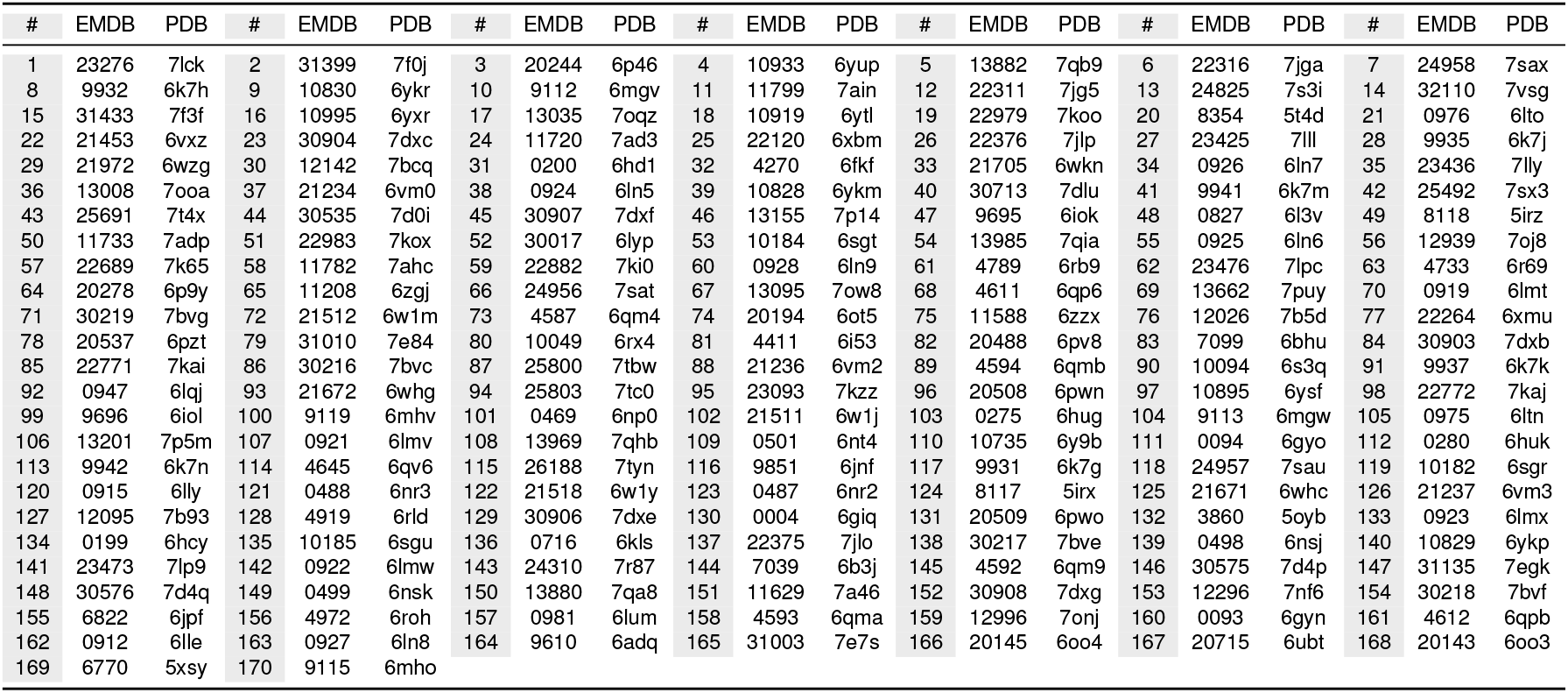
EMDB and PDB identifiers for maps and models used in SURFER training and validation set.

**Table S2.** EMDB and PDB identifiers for maps and models used in SURFER test set.

| # | EMDB | PDB | # | EMDB | PDB | # | EMDB | PDB | # | EMDB | PDB | # | EMDB | PDB | # | EMDB | PDB | # | EMDB | PDB |
| --- | --- | --- | --- | --- | --- | --- | --- | --- | --- | --- | --- | --- | --- | --- | --- | --- | --- | --- | --- | --- |
| 1 | 0257 | 6hra | 2 | 4032 | 5lc5 | 3 | 4272 | 6fki | 4 | 4288 | 6fo2 | 5 | 4588 | 6qm5 | 6 | 4589 | 6qm6 | 7 | 4646 | 6qvb |
| 8 | 4746 | 6r7x | 9 | 7009 | 6ave | 10 | 7133 | 6bqv | 11 | 7882 | 6dg7 | 12 | 8702 | 5vkq | 13 | 8958 | 6e1n | 14 | 8960 | 6e1p |
| 15 | 9934 | 6k7i | 16 | 9939 | 6k7l |  |  |  |  |  |  |  |  |  |  |  |  |  |  |  |

**Table S3.** SCUNet architecture summary.

| Layer (name) | Input | Output | InC | OutC | Params | Kernel |
| --- | --- | --- | --- | --- | --- | --- |
| SCUNet | 48<sup>3</sup> | 48<sup>3</sup> | 1 | 1 | — | — |
| Head | 48<sup>3</sup> | 48<sup>3</sup> | 1 | 32 | — | — |
| Conv3d | 48<sup>3</sup> | 48<sup>3</sup> | 1 | 32 | 864 | 3<sup>3</sup> |
| Down-sampling block 1 | 48<sup>3</sup> | 24<sup>3</sup> | 32 | 64 | — | — |
| ConvTransBlock (0) | 48<sup>3</sup> | 48<sup>3</sup> | 32 | 32 | 19,469 | — |
| ConvTransBlock (1) | 48<sup>3</sup> | 48<sup>3</sup> | 32 | 32 | 19,469 | — |
| Conv3d (2) | 48<sup>3</sup> | 24<sup>3</sup> | 32 | 64 | 16,384 | 2<sup>3</sup> |
| Down-sampling block 2 | 24<sup>3</sup> | 12<sup>3</sup> | 64 | 128 | — | — |
| ConvTransBlock (0) | 24<sup>3</sup> | 24<sup>3</sup> | 64 | 64 | 76,826 | — |
| ConvTransBlock (1) | 24<sup>3</sup> | 24<sup>3</sup> | 64 | 64 | 76,826 | — |
| Conv3d (2) | 24<sup>3</sup> | 12<sup>3</sup> | 64 | 128 | 65,536 | 2<sup>3</sup> |
| Down-sampling block 3 | 12<sup>3</sup> | 6<sup>3</sup> | 128 | 256 | — | — |
| ConvTransBlock (0) | 12<sup>3</sup> | 12<sup>3</sup> | 128 | 128 | 305,204 | — |
| ConvTransBlock (1) | 12<sup>3</sup> | 12<sup>3</sup> | 128 | 128 | 305,204 | — |
| Conv3d (2) | 12<sup>3</sup> | 6<sup>3</sup> | 128 | 256 | 262,144 | 2<sup>3</sup> |
| Bottleneck | 6<sup>3</sup> | 6<sup>3</sup> | 256 | 256 | — | — |
| ConvTransBlock (0) | 6<sup>3</sup> | 6<sup>3</sup> | 256 | 256 | 1,216,616 | — |
| ConvTransBlock (1) | 6<sup>3</sup> | 6<sup>3</sup> | 256 | 256 | 1,216,616 | — |
| Up-sampling block 3 | 6<sup>3</sup> | 12<sup>3</sup> | 256 | 128 | — | — |
| ConvTranspose3d (0) | 6<sup>3</sup> | 12<sup>3</sup> | 256 | 128 | 262,144 | 2<sup>3</sup> |
| ConvTransBlock (1) | 12<sup>3</sup> | 12<sup>3</sup> | 128 | 128 | 305,204 | — |
| ConvTransBlock (2) | 12<sup>3</sup> | 12<sup>3</sup> | 128 | 128 | 305,204 | — |
| Up-sampling block 2 | 12<sup>3</sup> | 24<sup>3</sup> | 128 | 64 | — | — |
| ConvTranspose3d (0) | 12<sup>3</sup> | 24<sup>3</sup> | 128 | 64 | 65,536 | 2<sup>3</sup> |
| ConvTransBlock (1) | 24<sup>3</sup> | 24<sup>3</sup> | 64 | 64 | 76,826 | — |
| ConvTransBlock (2) | 24<sup>3</sup> | 24<sup>3</sup> | 64 | 64 | 76,826 | — |
| Up-sampling block 1 | 24<sup>3</sup> | 48<sup>3</sup> | 64 | 32 | — | — |
| ConvTranspose3d (0) | 24<sup>3</sup> | 48<sup>3</sup> | 64 | 32 | 16,384 | 2<sup>3</sup> |
| ConvTransBlock (1) | 48<sup>3</sup> | 48<sup>3</sup> | 32 | 32 | 19,469 | — |
| ConvTransBlock (2) | 48<sup>3</sup> | 48<sup>3</sup> | 32 | 32 | 19,469 | — |
| Tail | 48<sup>3</sup> | 48<sup>3</sup> | 32 | 1 | — | — |
| Conv3d (0) | 48<sup>3</sup> | 48<sup>3</sup> | 32 | 1 | 864 | 3<sup>3</sup> |
| Total parameters: 4,729,084 |  |  |  |  |  | Trainable: 4,727,676 |
|  |  |  |  |  |  | Non-trainable: 1,408 |

**Fig. S1.**
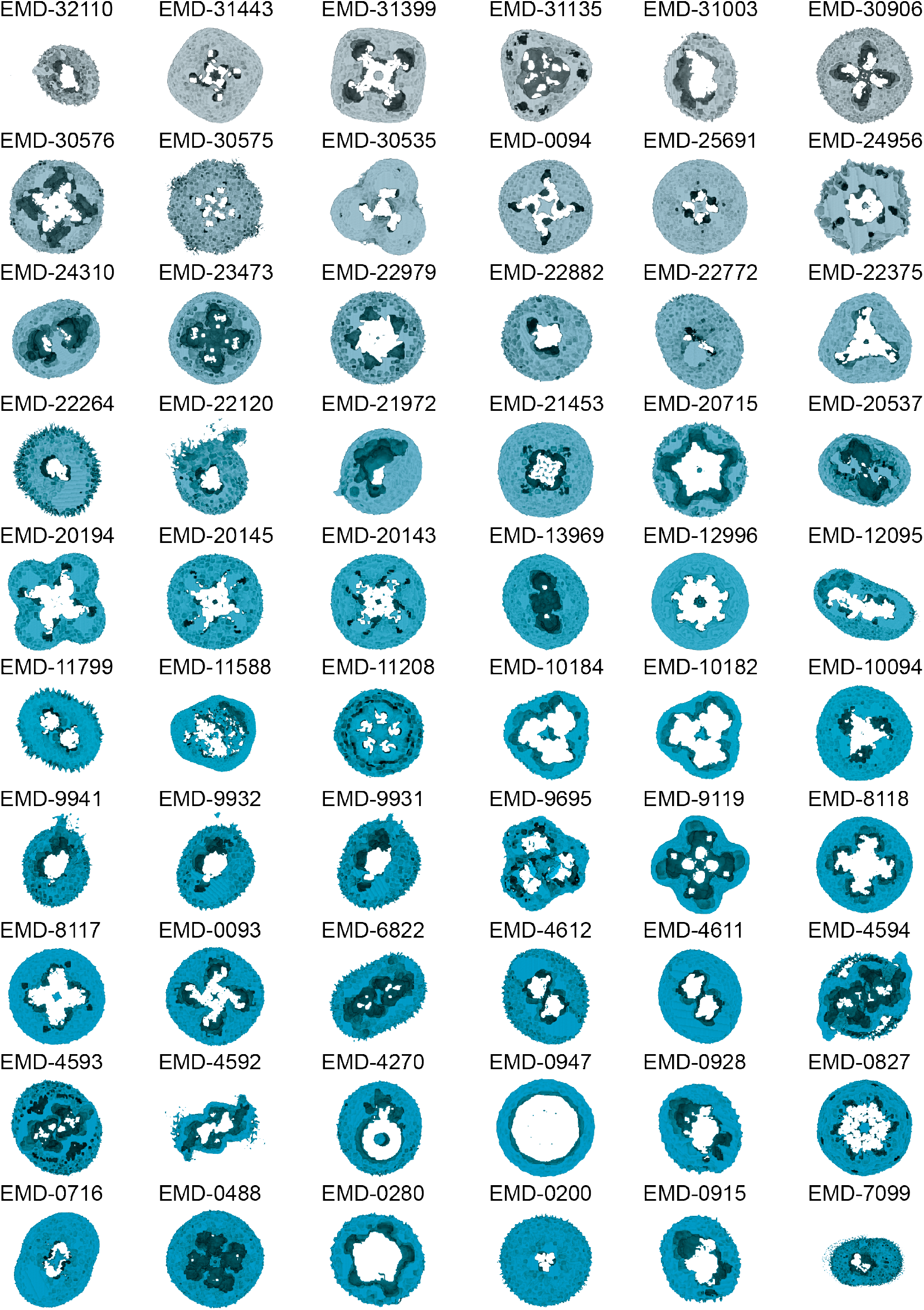
Shape variety of segmented membrane mimic densities. Randomly selected examples of segmented membrane mimic densities in the training data.

**Fig. S2.**
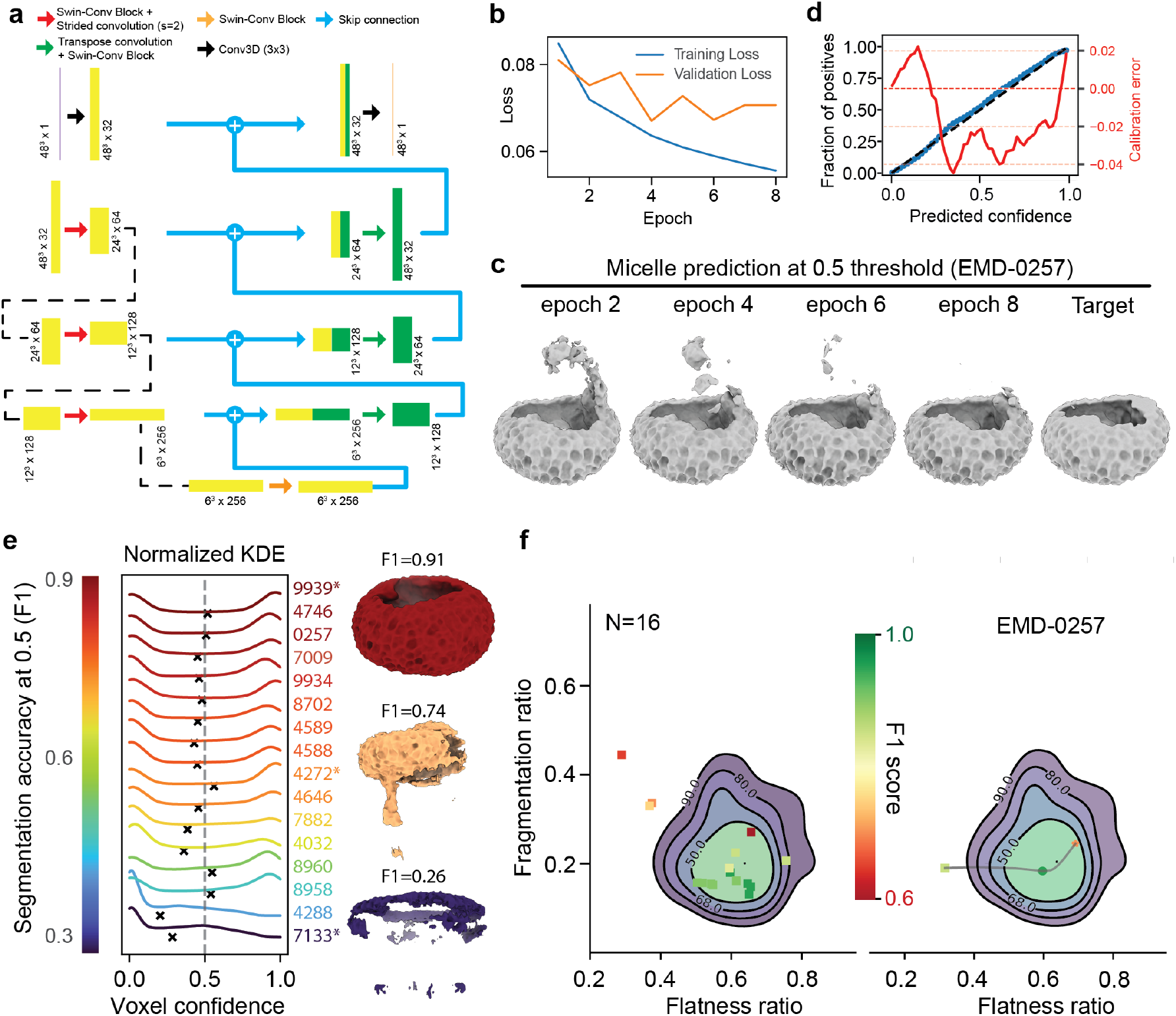
Network architecture, training behaviour and confidence characteristics of SURFER predictions.. (a) Network architecture of the SURFER SCU-Net. (b) Training and validation loss curves over eight epochs, showing stable optimisation without divergence between training and validation loss. (c) Predicted detergent micelle segmentation for a held-out test map (EMD-0257) using model checkpoints from successive epochs, binarised at a fixed threshold of 0.5. The overall micelle geometry is recovered early during training, while continued optimisation primarily reduces false-positive regions and sharpens the micelle boundary at the same threshold. (d) Reliability diagram aggregating voxel-wise predictions across all test maps, showing predicted confidence versus the observed fraction of true positives. The red curve indicates calibration error (predicted confidence minus observed fraction), with negative values corresponding to under-confidence. (e) Kernel density estimates of voxel-wise predicted confidence for individual test maps, weighted by the relative micelle volume within the molecular boundary. High-accuracy segmentations (top) show clearly bimodal confidence distributions with peaks near 0 and 1, whereas lower-accuracy cases (bottom) exhibit a skew toward low-confidence values, reflecting bias toward the majority class. Representative segmentations binarised at a threshold of 0.5 are shown alongside each distribution, with F1 scores indicated. (f) Left: Fragmentation and flatness ratios of the 16 test maps, thresholded at their respective optimal F1 scores, superimposed on the flatness–fragmentation distribution of the SURFER training set (see Figure 3c). Right: Trajectory of one representative test map (EMD-0257) across different binarisation thresholds (square: 0.1; circle: 0.5; asterisk: 0.9). Data points are coloured according to F1 score.

